# Disabling of ARC1 through CRISPR/CAS9 leads to a complete breakdown of self-incompatibility responses in *Brassica napus*

**DOI:** 10.1101/2022.08.09.503242

**Authors:** Kumar Abhinandan, Neil M.N. Hickerson, Xingguo Lan, Marcus A. Samuel

## Abstract

Self-incompatibility (SI) is a genetic mechanism utilized by many flowering plants to promote outcrossing by recognizing and rejecting self-pollen. While the molecular mechanisms controlling SI can differ in various families, plants in the Brassicaceae family have been found to regulate SI through a S-haplotype-specific receptor-ligand interaction. The recognition of self-pollen leads to the activation of ARMADILLO-REPEAT-CONTAINING1 (ARC1), an E3 ligase which is thought to degrade the compatibility factors required for pollen germination, thus resulting in SI. However, the role of ARC1 in SI in the Brassicaceae family has been disputed within the scientific community. While various studies demonstrate that the manipulation of ARC1 and its downstream targets can break-down the SI response, others show that the SI phenotype can be imparted to self-compatible cultivars without the ARC1 gene. This study investigated the involvement of ARC1 in SI through the generation of loss-of-function *ARC1* mutants using CRISPR-Cas9 in *B. napus.* Loss of *ARC1* resulted in complete breakdown of SI in two different S-haplotypes providing strong evidence for the essential nature of ARC1 for mounting a successful SI response.

## Introduction

Many flowering plants utilize self-incompatibility (SI) response as a genetic mechanism to prevent self-pollen from establishing on the stigmas to promote outcrossing and genetic diversity. In Brassica, during SI, recognition of the pollen ligand SP11 by S-locus receptor kinase (SRK) results in activation of an E3 ligase, Arm Repeat Containing protein (ARC1), that leads to proteasomal degradation of compatibility factors required for successful pollen acceptance. ARC1 was originally identified as an interactor of SRK kinase domain and is highly expressed in mature stigmas (Gu et al., 1998). Suppression of ARC1 through antisense technology resulted in partial breakdown of SI in the self-incompatible *Brassica napus* W1 line, establishing its role as a positive regulator of SI (Stone et al., 1999). This was further supported by the RNAi-mediated suppression of ARC1 in the self-incompatible *A. lyrata,* resulting in partial breakdown of the SI pathway (Indriolo et al., 2012). The observed partial compromise of SI in both Brassica and Arabidopsis suggested that either an alternative SI pathway or the incomplete suppression of ARC1 could have resulted in the incomplete breakdown of SI. This aspect has remained unresolved.

Despite the partial breakdown of SI shown in two different systems, the role of ARC1 in the Brassicaceae SI response has been questioned for the past couple of decades. Stable co-expression of *Brassica* SLG, SRK, and ARC1 was insufficient in conferring the SI phenotype in compatible *Arabidopsis thaliana* (Bi et al., 2000). In another report, when *A. thaliana* plants were transformed to express SRKb and SCRb genes, a strong SI phenotype was observed in the absence of ARC1 (Kitashiba et al., 2011). *A. thaliana* plants that exhibit SI phenotype through the transformation of SRK and SCR alone show similar trends of heritability, developmental regulation, and intensity of self-incompatibility response as self-incompatible cultivars of *Brassica* (Nasrallah and Nasrallah, 2014). Although, evolutionarily, Arabidopsis that lacked a bona fide ARC1 ortholog could have re-purposed other E3 ligases to assume the role of ARC1, the question of how complete loss of ARC1 could influence SI in Brassica sp. remained unresolved.

## Results

To unequivocally examine the role of ARC1 during SI, we decided to create *ARC1* loss-of-function *B. napus* by utilizing the CRISPR/Cas9 platform. In order to efficiently design specific targets for the CRISPR constructs, available gene copies of ARC1 or similar genes present in *B. napus* (allotetraploid) were retrieved (http://cbi.hzau.edu.cn/bnapus/index.php) (Song et al., 2020). After validation of the sequences across the published NCBI database, TOPO cloning was performed with the *BnARC1* specific cDNA amplification products obtained from the RNA pool of fully mature stigma of *B. napus.* Sanger sequencing of the clones revealed that *BnARC1* is a single copy gene derived from *Brassica rapa,* while its homolog from the C-genome was identified at low frequency and was most similar to PUB17 family genes (Fig. 1A). When the expression profiles of these two genes were assessed during various stages of stigma maturity, ARC1 had significantly higher expression than *PUB17* and peaked at stigma maturity (Fig. 1B).

**Figure 1.**
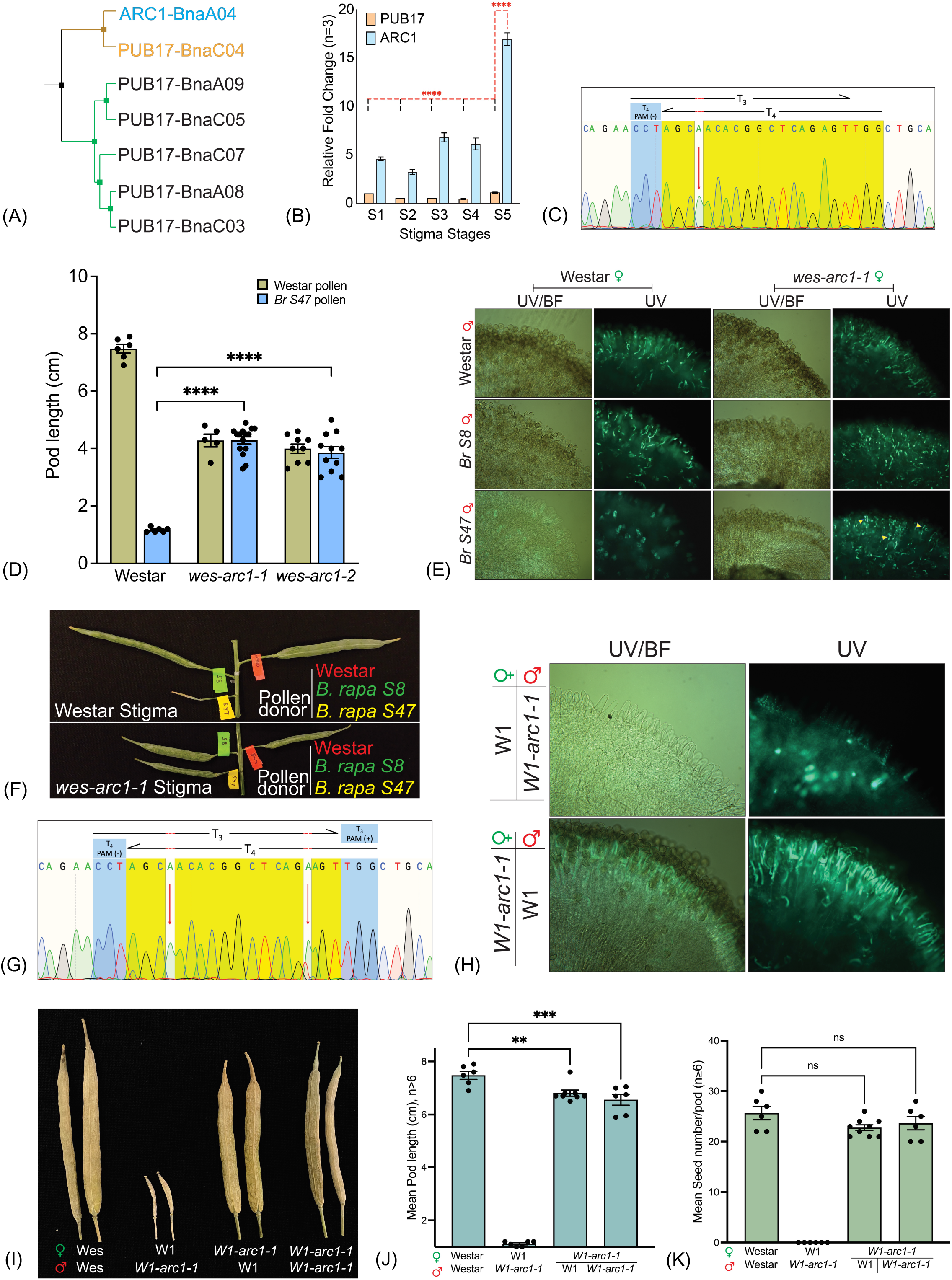
CRISPR–CAS9-mediated editing of ARM Repeat Containing 1 (ARC1) leads to a complete breakdown of Self-incompatibility response in *Brassica napus*. (A) ARC1 and related sequences were retrieved from the BnPIR *(Brassica napus* pangenome information resource) database. Highly divergent *PUB17 (PUB17-BnaC03a)* is not shown in the cladogram. (B) Relative expression of *ARC1* and *PUB17* during stigma development was assessed by qPCR. The bars show the fold change between *BnPUB17* and *BnARC1* during stigma development. Error bars represent the standard error of the mean (±SEM) (C) Chromatogram generated from Sanger sequencing of *BnARC1* from *wes-arc1-1*. Sequence spans the T3 and T4 target sites within the *BnARC1* coding region. The red arrow indicates the biallelic insertion of an ‘A’ at precisely 3 bases from the T4 protospacer adjacent motif (PAM) sequence indicating the insertion was created during an erroneous endogenous repair of the double stranded break performed by Cas9 at the T4 position. (D) Bar graph representing the average pod length in *wes-arc1* when pollinated with compatible Westar or incompatible *S47* pollen. The error bars represent the standard error of the mean (±SEM). (E) Aniline blue assay to detect pollen attachment and pollen tube penetration in flowers pollinated with pollen from Westar, S8 or *S47* haplotypes obtained from *B. rapa*, 24 h post-anthesis. The images were obtained either with UV (280-390 nm) to visualized aniline blue stain or with a green filter to observe pollen attachment. (F) Representative picture of pods still attached to inflorescence, pollinated with Westar (Self), *S47* (SI), and *S8* (CP) showing the breakdown of SI *wes-arc1-I* line. (G) Chromatogram generated from Sanger sequencing of *BnARC1* from *W1-arc1-1.* Sequence span the T3 and T4 target sites within the *BnARC1* coding region. The red arrows indicate the biallelic insertion of an ‘A’ at precisely 3 bases from the T4 PAM sequence and the biallelic insertions of an ‘A’ or ‘T’ at 3 bases from the T3 PAM sequence. (H) Aniline blue assay to assess pollen attachment and pollen tube penetration in reciprocal crosses of the CRISPR edited line *W1-arc1-1,* showing breakdown of SI in the absence of *ARC1.* (I) Representative picture of mature pods from hand-pollinated individual flowers with combinations shown. The full extension of pods after incompatible pollination indicates compromised SI in *W1-arc1-1* lines. (J, K) Bar graphs representing the average pod length (J) and seed set (K) in the *W1-arc1-1* when pollinated with W1 or self-pollinated comparable to Westar self-pollination. Error bars represent the standard error of the mean (±SEM).

We next created a multiplex CRISPR–Cas9 ARC1 editing construct and generated several *B. napus* Westar (SC) transgenic lines harboring the editing system through *Agrobacterium-mediated* transformation (Ma et al., 2015; Stanic et al., 2021). When *ARC1* was amplified from these transgenic lines, several of these lines possessed biallelic edits (Fig. 1C) (only T3/4 shown from *wes-arc1-1*).

The Westar cultivar is self-compatible due to an insertion in the promoter region of the SP11, leading to a lack of expression of SP11, the pollen ligand required for SI response (Okamoto et al., 2007). However, it retains all the downstream SI signaling components (SRK and ARC1) as it readily rejects pollen of *S47* haplotype origin with a functional SP11 (Okamoto et al., 2007). T2 generation of two independently edited *wes-arc1* plants with biallelic edits were subjected to pollination assays with compatible Westar or *S8* pollen and the incompatible *S47* pollen. When *B. rapa S47* haplotype pollen was applied to the control Westar stigma, a robust SI was observed in the Westar stigmas, which readily rejected pollen of the *S47* haplotype (Fig. 1D-F). In contrast, the *wes-arc1-1 and wes-arc1-2* plants showed a complete breakdown of SI response when pollinated with *S47* pollen (Fig. 1D - F). When either Westar pollen or *B. rapa* pollen from an *S8* haplotype (*SRK8* is absent in Westar) was used as a positive control for stigma receptivity, full acceptance was observed in both Westar controls and the *wes-arc1* plants indicating that stigma receptivity was not modified in the edited line (Fig. 1E, F). We also consistently observed that ARC1 defective plants produced pods that were shorter than the parental Westar plants suggesting that ARC1 should be involved in pod development, in addition to mediating SI response.

We next sought to test whether nullifying *ARC1* in the highly characterized self-incompatible W1 background would result in a complete breakdown of SI. The isogenic, self-incompatible W1 line was created by introgression of the dominant 910 *B. rapa* haplotype (SRK 910/SP11-910) and displays a very strong SI phenotype when self-pollinated but is compatible when crossed with Westar. We predicted that regardless of the upstream receptor/ligand complex, abolishing ARC1 function should lead to complete breakdown of SI.

Since W1 plants are incompatible and difficult to propagate/regenerate through conventional *Agrobacterium-based* tissue culture approaches, W1 plants were crossed with the Westar transgenic lines harboring the *ARC1* editing system. In the F1 generation, we identified a *W1-arc1-1* line which displayed strong breakdown of SI when self-pollinated. Sequencing of *ARC1* revealed biallelic edits (Fig. 1G, only T3/4 shown). When pollen from *W1-arc1-1* were tested on W1 stigmas, W1 stigmas rejected *W1-arc1-1* pollen indicating that *SP11-910* haplotype was not altered or deleted. On the other hand, *ARC1* edited *W1-arc1-1* plants readily accepted pollen from self-incompatible W1 plants (Fig. 1H). This complete breakdown resulted in a full seed set (Fig. 1I) that was comparable to when stigmas were pollinated with compatible Westar pollen (Fig. 1J and K).

## Discussion

We have convincingly demonstrated that elimination of ARC1 results in complete breakdown of SI in two different S-haplotypes *(S47* and *SRK910),* confirming the essential nature of ARC1 for SI response in *Brassica*. Under both situations, the closely related PUB17 ortholog was unaltered, indicating the exclusive nature of ARC1 for mediating SI response. Although this study eliminates any doubt whether ARC1 is required for SI in Brassica, the fact that in *A. thaliana,* ARC1 was shown to be dispensable in certain cases suggests that there could be an evolutionary significance to this observation. In self-incompatible *A. lyrata* ecotype, ARC1 is intact and required for SI response (Indriolo et al., 2012; Indriolo and Goring, 2014), while in self-compatible *Arabidopsis* species, *ARC1* is often found deleted (Indriolo et al., 2012; Indriolo et al., 2014). During the switch from SI to compatibility, *A. thaliana* could have either lost the *ARC1* gene and other components of the SI pathway or neo-functionalized them for various other pathways. Alternatively, ARC1 function could be species-specific as shown in a recent report, where the *A. thaliana* SC transgenic lines overexpressing SCR-SRK-ARC1 from *A. halleri* displayed SI phenotype, while they failed to manifest the SI phenotype when *ARC1* gene was derived from *Brassica napus*. These results indicate that the display of incompatibility in the SC *A. thaliana* might have genus-specific preferences (Zhang et al., 2019).

Nevertheless, this investigation has clearly demonstrated that absence of functional *BnARC1* in Westar and W1 plants leads to their inability to mount a successful SI showing that ARC1 is an indispensable downstream effector of SI in *Brassica.*

